# Diagnosis of type 2 Diabetes Mellitus (T2DM) using Paired microRNAs

**DOI:** 10.1101/2022.09.29.510072

**Authors:** Yukichi Takada, Yasuhiro Ono, Tatsuki Shibuta, Ayaka Ishibashi, Ayako Takamori, Kazuma Fujimoto, Yoshitaka Hirooka, Tsukuru Umemura

**Affiliations:** Department of Medical Technology and Sciences, Graduate School, International University of Health and Welfare, Okawa 831-8501, Japan; Diabetology, Kouhoukai Takagi Hospital, Okawa 831-8501, Japan; Department of Medical Technology and Sciences, International University of Health and Welfare, Okawa 831-8501, Japan; Clinical Laboratory, Kouhoukai Takagi Hospital, Okawa 831-8501, Japan; Medical Biostatistics, Saga University, Nabeshima 849-8501, Japan; Gastroenterology, Kouhoukai Takagi Hospital, Okawa 831-8501, Japan

## Abstract

Type 2 Diabetes mellitus (T2DM) is one of the most common diseases in the world and its prevalence ratio is still increasing. Patients with T2DM have diverse pathophysiological changes like as macrovascular, microvascular diseases, cancers as well as abnormal glucose metabolism. Thus, there are urgent needs to develop relevant biomarkers for the broad range of pathophysiology in patients with T2DM. We analyzed the signatures of serum miRNAs with the miRNA array analysis and reverse-transcription based quantitative polymerase chain reaction (RT-qPCR) in 50 patients with type 2 DM (T2DM) and 15 normal subjects. Array analysis showed that 19 miRNAs were up-regulated more than 2-fold and 71 miRNAs were down-regulated less than 0.5 in T2DM in comparison with normal subjects. Top 5 of up-regulated miRNAs were miR-3619-3p, miR-557, miR-6850-5p, miR-3648, miR-4730, and 5 of most down-regulated miRNAs were miR-5100, miR-4454, miR-1260b, miR-7975, miR-6131. We selected 4 miRNAs for validation analysis with RT-qPCR based on the abundance enough for reliable analyses and disease-specificities reported in previous reports. Serum miR-126-3p was down-regulated (3.21-fold, p<0.05) in T2DM, and miR-10a up-regulated (1.94-fold, p<0.05). However, none of single miRNA had significant correlation with clinical data and state. Data of the paired miRNAs: miR-10a and miR-200c, or miR-126 and miR-10a, clearly differentiated T2DM patients from normal subjects (p<0.05). Our study showed the paired-miRNA analyses as the more effective diagnostics for T2DM than the single miRNA analysis.

## Introduction

Type 2 Diabetes mellitus (T2DM) is one of the most common diseases in the world and its prevalence ratio is still increasing [1,2]. Diabetes mellitus is categorized into two types [1,3]. Type 1 DM (T1DM) is characterized with dysfunction of pancreatic β cells caused by immunological mechanism and insulin deficiency [4,5]. Type 2 DM (T2DM) develops by insulin resistance or insulin deficiency with background of several risk factors: genetic, obesity, deficiency of exercise, stress etc. [1-3]. Almost 95% of DM patients is T2DM, of whom population is over 400 million in the world [1,3]. Diagnosis of T2DM depends on abnormalities of glucose metabolisms indicated by blood sugar and glycated hemoglobin A1c (HbA1c) levels [1,6]. About 10% of T2DM was presented with diabetic ketoacidosis at diagnosis [2]. However, mortality and quality of life (QOL) of patients with T2DM depends on complicated cardiovascular disease (CVD) [7-10] or chronic kidney disease (CKD) [8,11] [12,13]. CVD was diagnosed in 32.2% of patients with T2DM and was the cause of death in 9.9% in patients with T2DM [10]. Incidence of heart failure of T2DM patients is 0.5% which is almost three times higher than in healthy population [3].Kidney disease was present 42.3% of patients with T2DM while 9.4% of non-T2DM persons and was the important cause of increased mortality in T2DM [13]. Quality of life (QOL) of T2DM patients is also disturbed by other complications than CVD or KD, namely infectious diseases, cancers, retinopathy, or neuropathy [1,2,14,15]. Therefore, biomarkers to detect the wide range of pathophysiological changes may lead us to precision medicine of each diabetic patient.

MicroRNAs (miRNAs) are small non-coding RNAs which regulates gene expression at the posttranscriptional stage by digesting target messenger RNAs (mRNAs) or inhibiting the translation step [16,17]. miRNAs are involved in development, cellular proliferation and differentiation, apoptosis, and tumor genesis [18,19]. Some parts of miRNAs are released into extracellular space or circulating blood. Extracellular vesicles (EVs) are nano-sized particles (50 to 1,000 nm) to transport miRNAs to recipient cells at the distance organs [20,21]. Thus, circulating miRNAs are important molecules for cell-to-cell communication. Dysregulation of miRNAs was reported as a possible mechanism of diabetic neuropathy and nephropathy [22-27]. In this study, we focus on circulating microRNAs as relevant biomarkers for early diagnosis and systemic complications of patients with T2DM.

## Material and Methods

### Patients

In this study we recruited 50 patients with T2DM and 15 control subjects in Kouhoukai Takagi Hospital, Ohkawa, Japan or International University of Health and Welfare. Diagnosis and clinical staging of patients with T2DM were done according to criteria by Japanese Diabetic Society [28]. Control subjects included 9 males and 6 females whose ages were 21- to 56-year-old, without T2DM and noteworthy diseases. This study was approved by Research Ethics Committee in International University of Health and Welfare and Takagi Hospital, and informed consent was obtained from all subjects with the signed agreement form.

### Blood sampling and preservation

Serum samples were obtained after blood coagulation at 25 °C for 30 min, by centrifugation of 3,500 rpm for 10 min at room temperature. After passing membrane filter with 0.45 μm pore size, samples were kept in freezers at -80 °C until use [29,30].

### RNA isolation

Isolation of miRNAs from 300 μL of each serum sample was done with NucleoSpin miRNA Plasma (Takara Bio, Inc., Tsu, Japan) according to manufacturer’s instruction. Briefly, after adding 90 μL of lysis buffer to serum, serum was vortexed for 5 seconds and was kept in the tube stand for 3 min. Next, 30 μL of protein denaturant was added and the tube was vortexed for 5 seconds. Supernatant was obtained by centrifuging serum mixture at 11000 rpm for 3 min to remove protein precipitate. Supernatant of 300 μL was mixed with 400 μL isopropanol and 3 μL of *Caenorhabditis elegans* miR-39 (cel-miR-39) solution (1 fmol/μL) as a spike-in control in the new tube. Mixture of 700 μL was transferred into the column set on the new tube and was passed through the column at 11000 rpm for 30 seconds to catch nucleic acid on membrane. In the next steps, DNAs on the column were removed with DNase treatment and washing buffers. Finally, 30μL RNase-free-water was added onto the column and RNA solution was obtained by centrifuging at 11000 rpm for 1 minutes. RNA specimens were kept in -80°C in freezer [29,30].

### miRNA microarray analysis

For analysis of serum miRNA signatures 5 serum samples were pooled from healthy subject and T2DM patients whose HbA1c levels were 8.4-9.7%. miRNA array analyses were done using two pooled samples with DNA Chip 3D-Gene (Toray Industries, Inc., Japan). Each 3D-Gene Chip has DNA probes for over 2632 miRNAs and quantitatively detects miRNAs in samples [31].

### Quantitative analysis of serum miRNAs with RT-qPCR

Complementary DNAs (cDNAs) were made from extracted miRNAs with High-Capacity cDNA Reverse Transcription Kit (Thermo Fisher Scientific). Briefly, master mix was made with 0.1 μL dNTPs (100mM), 1 μL MultiScribe Reverse Transcriptase (50U/μL), 1.5 μL 10X RT buffer and 4.225 μL RNase-free-water for each sample. Seven μL of master mix was, 3 μL of target miRNA-specific primer and 5 μL cDNAs solution were added to the new tube. Reverse transcription reaction was done using the thermal cycler (Thermo Fisher Scientific.). The reaction condition was 16°C for 30 min, 42°C for 30 min and 85°C for 5 min. cDNAs were preserved in -20°C freezer. Quantitative polymerase chain reaction (qPCR) was done using TaqMan™ Fast Advanced Master Mix (Thermo Fisher Scientific) and the real time-PCR machine (ABI7500fast, Thermo Fisher Scientific). Briefly, mix 10μL 2X TaqMan™ Fast Advanced Master Mix, and 7.67 μL RNase-free-water in tube by 1 sample to make Master mix. Next, mix 17.67 μL Master mix, 1 μL primer according to target miRNA and 1.33 μL template made in RT step in new tube and spin down after tapping. The qPCR condition was 95°C for 10 mints, and 40 cycles of 95°C in 15 sec, 60°C for 60 sec [29,32].

### MiRNA analysis by 2*-ΔΔCT* Method

The 2*-ΔΔCT* Method is a major method of quantitate miRNA analysis to compare specific miRNAs at healthy and T2DM. Threshold Cycle (CT) value of each sample was defined as the number of minimum cycle number to show detectable fluorescent signal. To compare miRNA levels between healthy subjects and T2DM patients, difference of CT values (ΔCT) was calculated by subtracting CT values of target miRNAs by CT value of spiked-in cel-miR-39 (1 fmol/μL) at first and compare after further subtraction between values of healthy subjects and T2DM patients (ΔΔCT) [32].

### Statistical analysis

Significant test by Mann–Whitney U test: For statistical comparisons of healthy and T2DM, use Mann–Whitney U test which common nonparametric test in two groups. miRNA usually not depend on constant distribution so more appropriate than t-test which common parametric test in two groups. Correlation coefficient by Pearson’s correlation coefficient: Pearson’s correlation coefficient is common test to find correlation strength in two groups. In this study, use to finds correlation in multiple miRNAs in healthy and T2DM. Separation two groups by Logistic regression analysis: Logistic regression analysis is major test to separate two groups and useful to predict a probability of random sample belong to one group or not at scatter plot. In this study, use to statistical separate healthy and T2DM.

## Results

### Clinical data of patients with T2DM

Fifty patients with T2DM (34 males and 16 females) were studied (Table 1). Average of ages was 65.32 year (range 32 to 93). The mean of body mass index (BMI) was 25.7 (17.4 to 39.4). Indexes of glucose metabolism were all high: fasting glucose 168.9 mg/dL (76 to 364), HbA1c 7.8% (14.6 to 36.5), glycosylated albumin (GA) 21.1% (14.6 to 36.5). Thirty-two of 50 patients (64%) had neuropathy, 18 (36%) retinopathy, 34 (68%) nephropathy, according to criteria by Japanese Society of Diabetes, Japanese Society of Ophtalmic Diabetology (S2 Table) [33].

**Table 1.** Clinical data of patients with T2DM (n=50)

| Table 1 Clinical data of patients with T2DM (n=50) |  |  |
| --- | --- | --- |
| Indexes |  | data |
| Age |  | 65.3 (32-93) year old |
| Gender | male | 34 (68%) |
|  | female | 16 (32%) |
| BMI* |  | 25.7 (17.4-39.4) |
| fasting blood glucose |  | 168.9 (76-364) mg/dL |
| HbA1c |  | 7.8 (5.3-11.0) % |
| GA |  | 21.1 (15.6-36.5) % |
| Complications |  |  |
| Nephropathy |  | 34/50 (68%) |
| Retinopathy |  | 18/50 (36%) |
| Neuropathy |  | 32/50(64%) |
| Insulin therapy |  | 22/50(44%) |
| BMI: body mass index, HbA1c: glycated hemoglobin<br>A1c, GA: glycosylated albumin, |  |  |

### Microarray analysis of circulating miRNAs

To characterize the T2DM-specific signatures of serum miRNAs, the array analysis was done by using pooled serum from control subjects (n=5) and patients with T2DM (n=5). In total, 2632 miRNAs were analyzed with Human miRNA Oligo chip (3D Gene). Scatter plotting is shown in Fig 1. Ratios of miRNA levels between control subjects and T2DM patients were calculated using globally normalized values of higher than 100: the average of globally normalized level was 25. Nineteen miRNAs were up-regulated more than 2-fold and 71 miRNAs were down-regulated less than 0.5 of which top 20 highly down-regulated were shown in Table 2. Top 5 of up-regulated miRNAs were miR-3619-3p, miR-557, miR-6850-5p, miR-3648, miR-4730, and top 5 of down-regulated miRNAs were miR-5100, miR-4454, miR-1260b, miR-7975, miR-6131.

**Table 2.** Up- and down-regulated serum miRNAs in patients with T2DM: 19 up-regulated miRNAs with ratios over 2.0 and 20 of 71 downregulated miRNAs with ratios less than 0.5 were listed. Ratios were calculated by dividing the value in T2DM with the value in normal control obtained from the array analysis.

| Up-regulated |  |  | Down-regulated |  |  |
| --- | --- | --- | --- | --- | --- |
| rank | miRNA | ratio | rank | miRNA | ratio |
| 1 | hsa-miR-3619-3p | 7.665 | 1 | hsa-miR-5100 | 0.086 |
| 2 | hsa-miR-557 | 5.972 | 2 | hsa-miR-4454 | 0.117 |
| 3 | hsa-miR-6850-5p | 4.210 | 3 | hsa-miR-1260b | 0.137 |
| 4 | hsa-miR-3648 | 4.080 | 4 | hsa-miR-7975 | 0.138 |
| 5 | hsa-miR-4730 | 3.944 | 5 | hsa-miR-6131 | 0.150 |
| 6 | hsa-miR-4745-5p | 3.808 | 6 | hsa-miR-4688 | 0.157 |
| 7 | hsa-miR-642b-3p | 3.725 | 7 | hsa-miR-744-5p | 0.177 |
| 8 | hsa-miR-937-5p | 3.696 | 8 | hsa-miR-7977 | 0.189 |
| 9 | hsa-miR-642a-3p | 2.934 | 9 | hsa-miR-4530 | 0.190 |
| 10 | hsa-miR-6826-5p | 2.813 | 10 | hsa-miR-6132 | 0.190 |
| 11 | hsa-miR-92a-2-5p | 2.698 | 11 | hsa-miR-4467 | 0.195 |
| 12 | hsa-miR-128-2-5p | 2.611 | 12 | hsa-miR-4505 | 0.201 |
| 13 | hsa-miR-4649-5p | 2.357 | 13 | hsa-miR-3679-5p | 0.201 |
| 14 | hsa-miR-6807-5p | 2.272 | 14 | hsa-miR-5001-5p | 0.207 |
| 15 | hsa-miR-8063 | 2.244 | 15 | hsa-miR-6820-5p | 0.214 |
| 16 | hsa-miR-6756-5p | 2.173 | 16 | hsa-miR-4443 | 0.216 |
| 17 | hsa-miR-6766-5p | 2.097 | 17 | hsa-miR-6126 | 0.229 |
| 18 | hsa-miR-10396a-3p | 2.076 | 18 | hsa-miR-10400-3p | 0.233 |
| 19 | hsa-miR-3663-3p | 2.015 | 19 | hsa-miR-10392-5p | 0.246 |
|  |  |  | 20 | hsa-miR-6893-5p | 0.253 |

**Fig 1.**
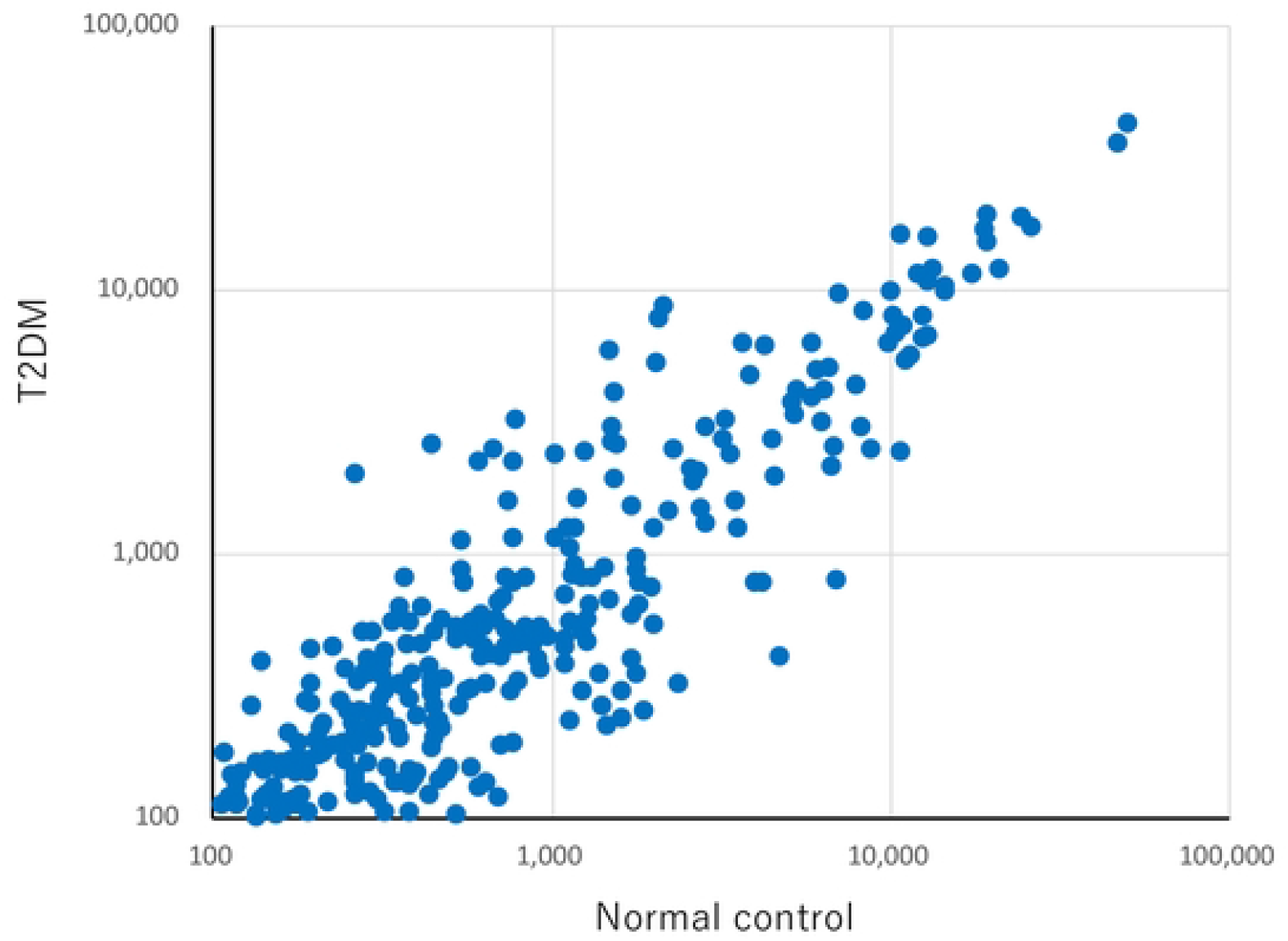
Most specific miRNAs at T2DM in RNA microarray. Results of array analysis of 2632 miRNAs were shown. Each dot is the normalized value using global average of normal control (x axis) and T2DM patients (y axis).

### Quantitative analysis of serum miRNAs in T2DM with RT-qPCR method

For validation study on dysregulated serum miRNAs in T2DM, we selected 4 miRNAs (miR-126-3p, miR-21, miR-10a, miR-200c) according to data from the array analysis and previous studies reporting relationship to T2DM clinical staging. Array data detected miR-126-3p in normal subject as 320th rank of 2632 miRNAs. Data of miR-21, miR-10a, miR-200) were not detected.

The level of serum miR-126 significantly decreased in T2DM group (1.96 ± 2.27, n=50) than healthy group (6.17 ± 1.85, n=50) (p<0.01) (Fig. 2A) and AUC is 0.98 (Fig. 2B). Best cut off level is 3.56 (sensitivity is 0.92 and specificity is 0.94). miR-10a significantly increased in T2DM group than healthy group (p<0.05) and AUC is 0.74. Best cut off level is 0.012 (sensitivity is 0.52 and specificity is 1.00). miR-21 and miR-200c did not show the significance differences between control subjects and T2DM patients (Figure 2A).

**Fig 2.**
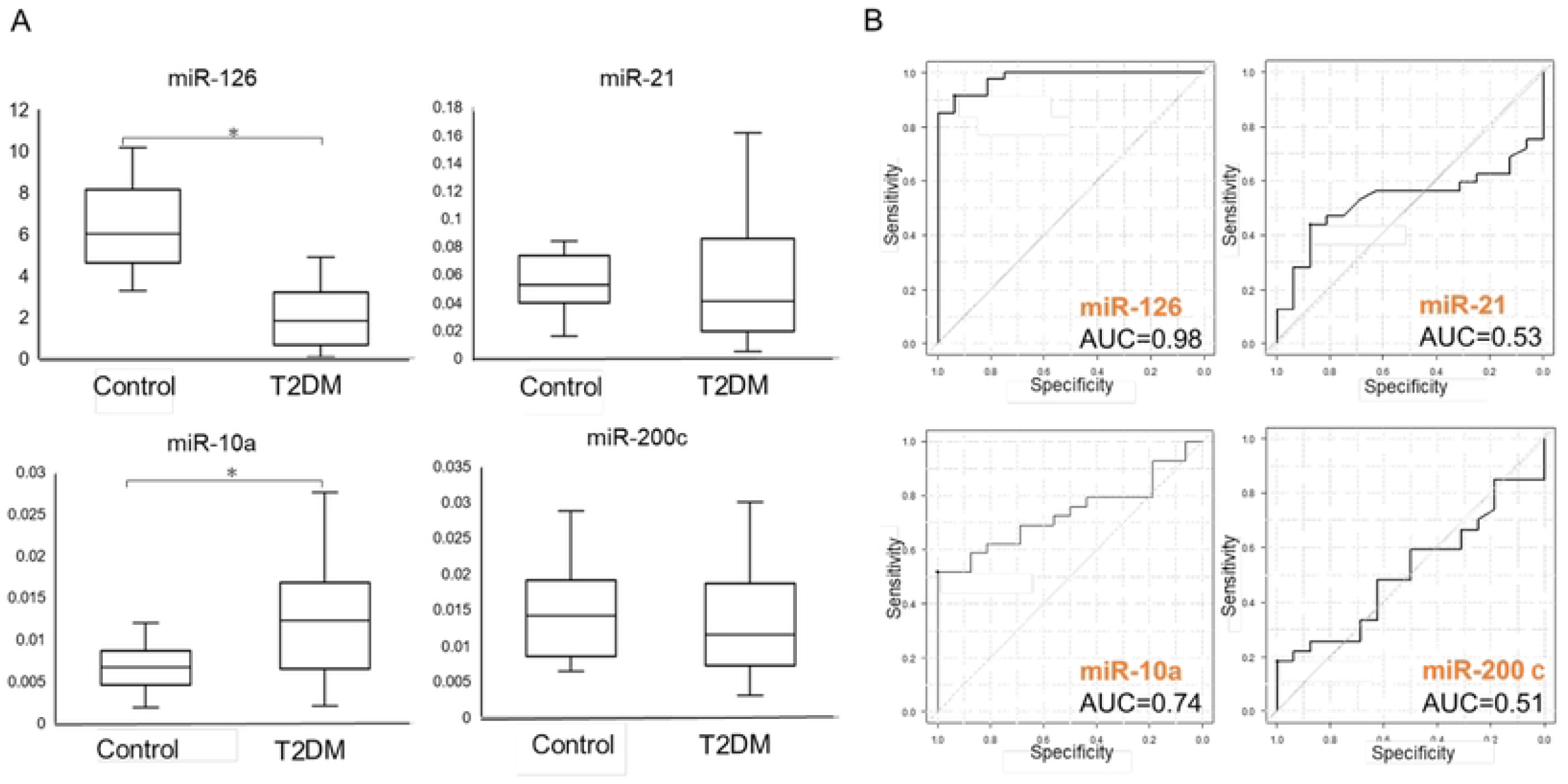
Serum levels and ROC analysis of four miRNAs. A. Box plot graphs of serum miRNAs levels analyzed with RT-qPCR. The y axis shows miRNA levels in control subjects (Control, n=15) and in T2DM patients (T2DM, n=50). Each result was normalized with the level of 1 femtomole cel-miR-39. Asterisk means significant difference between control subjects and T2DM patients (p<0.05). B. ROC curve and Area Under the Curve (AUC) of four miRNAs.

### The levels of miRNAs in different clinical stages of T2DM

The level of serum miR-126 was down- and that of miR-10c was up-regulated in 50 patients with T2DM. The levels of two miRNAs did not show any significant correlation with clinical data including HbA1c and GS (Table 3). Therefore, we evaluated two miRNAs in the different clinical stages. Clinical staging was done using HbA1c and GA according to criteria by Japanese Diabetic Society. Out of 50 patients, 9 patients had HbA1c levels less than 6.9 mg/dL, 20 patients, 7.0 to 7.9, 21 patients, over 8.8, respectively. Serum miR-126 significantly increased in parallel with HbA1c levels higher than 7.0 (p<0.05), although the levels were still lower than normal subjects (Fig 3A). miR-10a increased only in moderate or severe categories of T2DM, with higher HbA1 levels more than 7.0% (Fig 3A). In the staging by GA values, only miR-10a showed a good correlation with GA levels (Fig 3B). The levels of miR-21 and miR-200c did not have any significance correlation between HbA1c and GA staging.

**Table 3.** Correlation between miRNA levels and clinical data

| indexes | HbA1c | GA | miR-126 | miR-21 | miR-10a | miR-200c |
| --- | --- | --- | --- | --- | --- | --- |
| Age | 0.42* | 0.49* | -0.05 | 0.21 | 0.26 | -0.30 |
| BMI | 0.01 | -0.24 | -0.01 | -0.01 | -0.18 | 0.05 |
| Disease duraion | 0.40* | 0.22 | 0.07 | -0.12 | 0.12 | -0.10 |
| fasting glucose | 0.36 | 0.61* | 0.16 | 0.31 | 0.15 | 0.10 |
| HbA1c | 1.00 | 0.72* | 0.29 | 0.28 | 0.28 | 0.18 |
| GA | 0.72* | 1.00 | 0.30 | 0.22 | 0.34 | -0.01 |
| eGFR | 0.11 | -0.24 | -0.06 | -0.05 | -0.25 | 0.12 |
| urinary protein | -0.12 | 0.06 | 0.02 | 0.01 | 0.00 | -0.10 |
| urinary albumin | 0.14 | 0.25 | -0.11 | 0.01 | 0.00 | -0.08 |
| creatinin | -0.22 | 0.04 | 0.10 | 0.00 | 0.02 | -0.11 |
| BUN | -0.19 | 0.07 | 0.10 | -0.03 | 0.05 | -0.17 |
| TC | -0.04 | -0.06 | 0.02 | -0.08 | 0.05 | -0.01 |
| HDL-C | -0.16 | -0.10 | 0.00 | -0.20 | 0.01 | -0.13 |
| non-HDL-C | 0.01 | -0.03 | 0.02 | -0.01 | 0.05 | 0.05 |
| TG | 0.11 | 0.10 | 0.10 | 0.19 | -0.06 | 0.08 |
| ALT | -0.02 | -0.23 | -0.15 | 0.19 | -0.17 | -0.14 |
| AST | 0.03 | -0.11 | -0.14 | 0.24 | -0.10 | -0.21 |
| γ-GT | -0.05 | -0.15 | -0.15 | 0.03 | 0.09 | -0.04 |
| amylase | -0.32 | -0.15 | -0.19 | 0.04 | -0.03 | -0.11 |

\*: R value >0.4, #: eGFR, estimated glomerular filtration rate; BUN, blood urea nitrogen; TC, total cholesterol; C, chST, aspartate aminotransferase; ALT, alanine aminotransferase; γ-GT, gamma-glutamyl transpeptidase.

**Fig 3.**
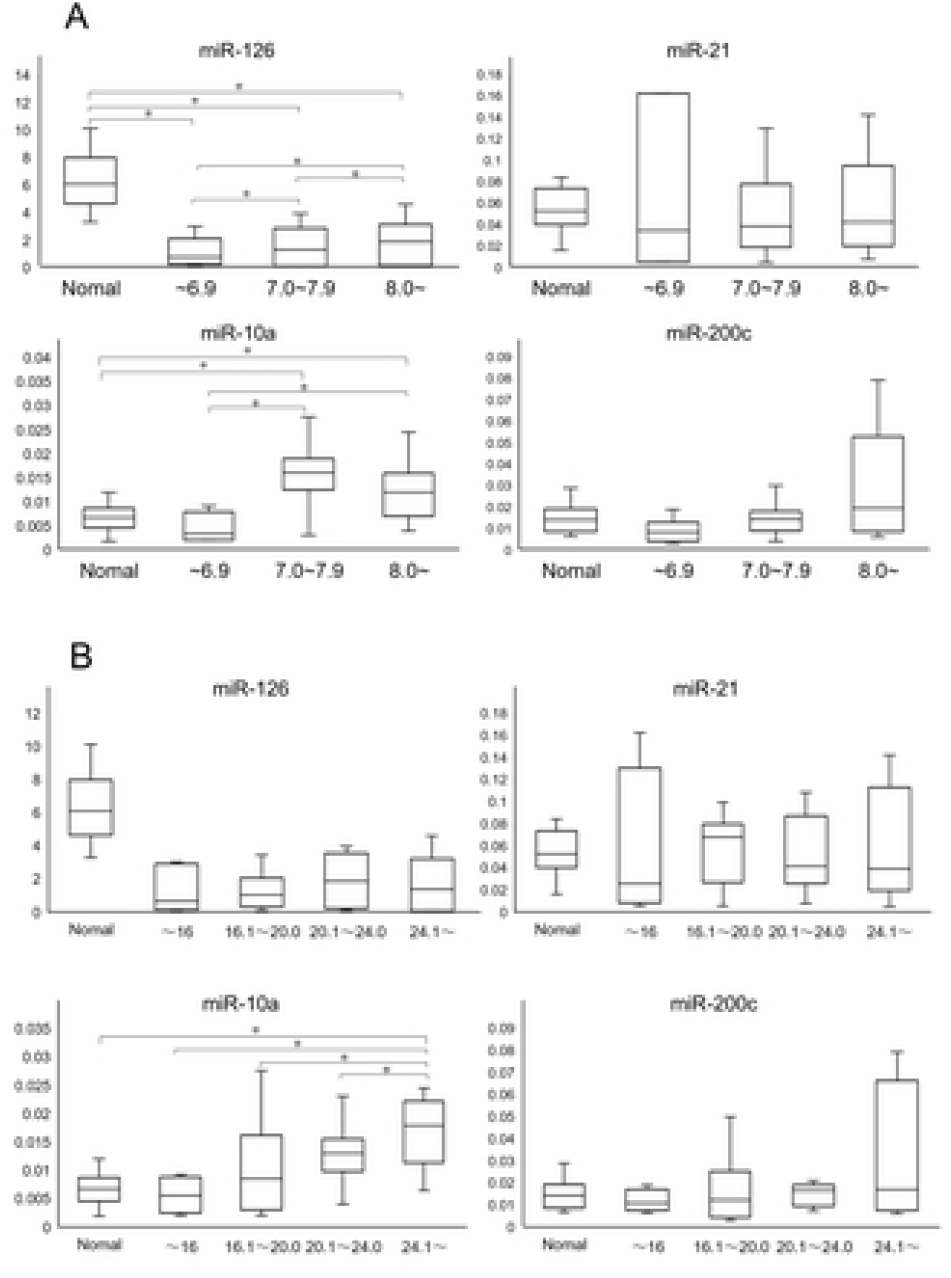
The levels of 4 MiRNAs according to the HbA1c and GA staging. The vertical axis shows data by box plot analysis of 4 miRNAs normalized with cel-miR-39. The horizontal axis shows the different stages according to HbA1c (A) and GA (B) in 50 T2DM patients (n=30 in case of miR-10a and miR-200c). The asterisks show significant differences between each stage (p<0.05). The recommended levels were under 7.0 (A) for HbA1c and under 16.0 for GA (B).

### Paired miRNA data is a useful biomarker for diagnosis of T2DM

Two miRNAs: miR-126 and miR-10a were dysregulated in patients with T2DM. None of each clinical data showed direct correlation with the levels of miR-126 or miR-10a. miR-21 and miR-200c did not show significant change in serum levels. (Table 3).

On the other hands, our data showed that patients with T2DM and normal subjects were separately placed on the scatter plot analysis using paired miRNAs: miR-21 vs miR-126, miR-21 vs miR-10a, miR-10a vs miR-126, miR-126 vs miR-200c (Fig 4, A, B, D and F). Logistic regression analysis showed the statistical significance of these separation of two groups (p<0.05). In cases of miR-21 vs miR-200c, miR-10a vs miR-200c, plots for two groups were not separated (Fig 4, C and E, respectively).

**Fig 4.**
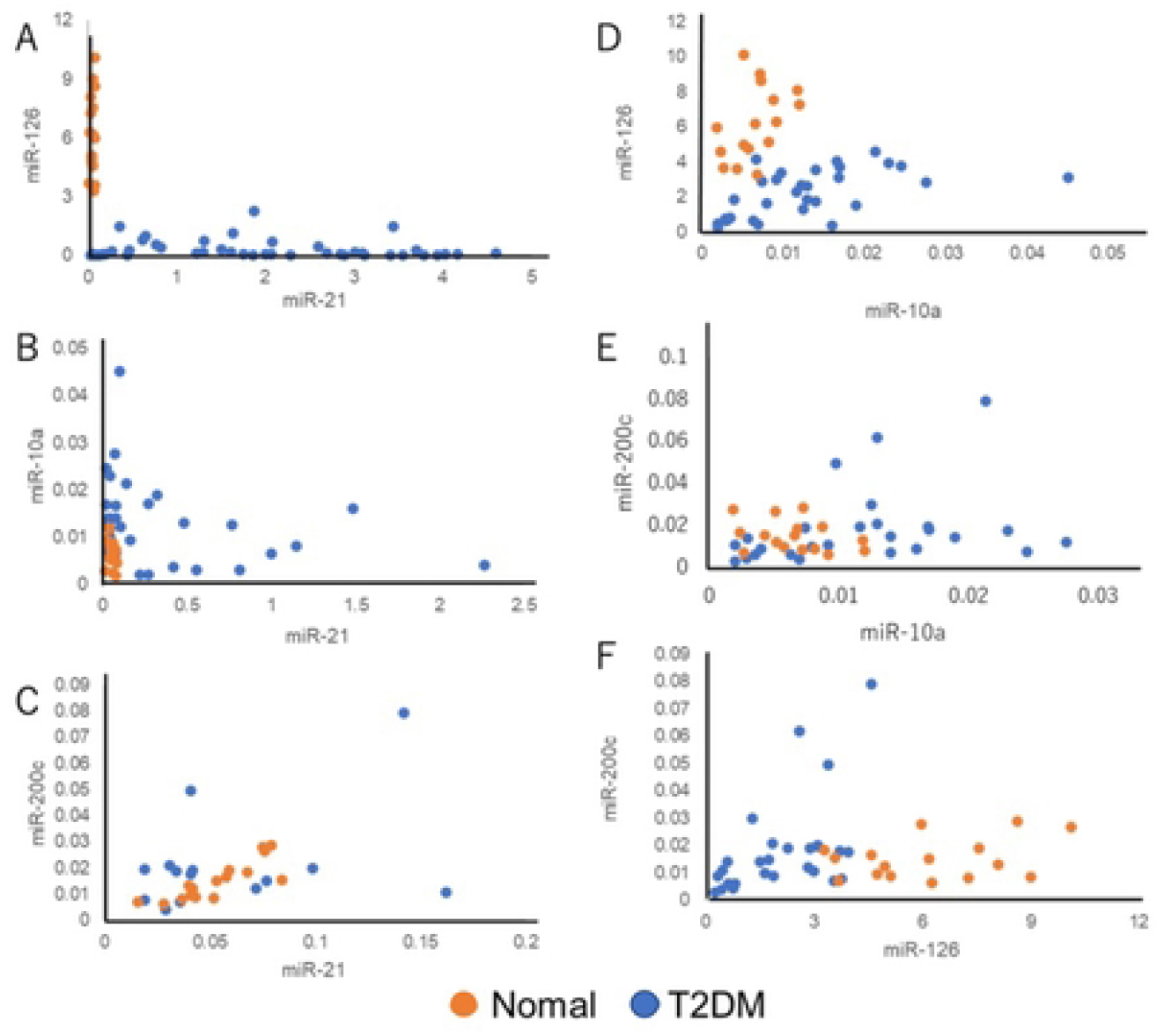
Differential pattern of paired miRNAs analysis between T2DM and normal control. The vertical axis and transverse axis in scatter plot show the levels of each miRNA normalized with cel-miR-39 in 50 T2DM patients (n=30 in case of miR-10a and miR-200c).

## Discussion

In this study, we tested the usefulness of circulating miRNAs signatures for diagnosis and staging of diseases. Microarray analysis of miRNAs using pooled serum from 5 patients with high HbA1c group of T2DM, showed 19 miRNAs up-regulated more than 2-fold in comparison with pooled sample from normal subjects. The functions of miRNAs in top rank were reported as important for cancer biology or cellular proliferation mainly examined in non-T2DM diseases or mouse models, for example, miR-3619-3p in papillary thyroid carcinoma and small cell carcinoma of the esophagus [34,35], miR-557 in osteosarcoma and pancreatic ductal adenocarcinoma [36,37], miR-6850 in serous ovarian carcinoma [38]. On the other hands, dysregulation of miRNAs in patients with T2DM have been reported: up-regulation of miR-100 and down-regulation of miR-126-3p [39,40]. Together with data of our array analysis and previous reports on miRNA dysregulation in T2DM, we selected 4 miRNAs (miR-126, miR-21, miR-10a and miR-200c) to analyze with RT-qPCR method in 50 patients with T2DM.

Our study showed that serum miR-126 was down-regulated in patients with T2DM. The same finding of down-regulation of miR-126 was reported in diabetic patients [41]. miR-126 is abundant in platelet and vascular endothelium and is involved in the mechanism to activate phagocytic activity of macrophages [42]. Down-regulation of miR-126 leads high risk of coronary artery disease [43]. In this our study, 36% of patients had retinopathy and 64% had neuropathy, suggesting complicated with diabetic microangiopathies (Table 1). miR-126 increased in kidney disease or myelodysplastic syndrome (MDS), suggesting the similar mechanism might up-regulate miR-126 at different stages of T2DM [44,45]. miR-10a known as marker of renal disorder [46] and increase in tubular damage in nephropathy [47]. Nephropathy observed in 68% of patients in this study might be reason of up-regulation of miR-10a. Taken together, dysregulation of miR-126 and miR-10a might reflect vascular damages in patients with T2DM.

There was no correlation between serum miR-126 and other indexes for T2DM staging. Therefore, miR-126 may be an independent marker for T2DM. Our data showed relative higher levels in advanced stages of T2DM. Thus, miR-126 might be a useful biomarker to predict vascular complications in T2DM patients. However, the higher levels of serum miR-126 in older people were reported [48]. Larger cohort is recommended to exclude this possibility that up-regulated levels of miR-126 might reflect older patients in advanced stages of T2DM. The levels of miR-10a were in parallel with the levels of HbA1c or GA in advanced stages of T2DM (Fig.3 A, B). miR-10a is one of miRNAs to promote vascular hyperactivity and organ damages in rats with diabetes and hyperlipidemia [49]. Our data did not have significant correlation with indexes of glucose or lipid metabolism. Expression of miR-10a was down-regulated in endothelial cells located at atherosclerosis-susceptible aorta than another portions in swine [50]. miR-10a in endothelial cell-derived EVs is transferred to monocytes and represses inflammatory signaling by inhibiting NF-κB pathway [51]. Thus, Up-regulation of serum miR-10a in T2DM with stage-dependent manner, may suggest protective response of human body against vascular damages in T2DM. MiR-21 is one of most important miRNAs in cancer biology [52-54]. Furthermore, many studies reported dysregulation of miR-21 in various diseases. Patients with T2DM have macro and microangiopathies with wide range of complications [2], which might result in no significant difference of miR-21 between T2DM and control subjects [52-57]. Several studies reported that miR-200c contributed to development of nephropathy in T2DM [58-60] and to lung cancer [61]. In our study, miR-200c expression level showed no significant difference in T2DM.

In this study we focused on development of the simple and efficient test of T2DM by using serum miRNAs profiles. Ochiya’s group showed high sensitivity and specificity of combined analysis of multiple serum miRNAs for detecting early cancers [62]. However, the similar approach was not done in T2DM. The analysis of paired miRNAs in our study showed useful differentiation between patients with T2DM: miR-21 vs miR-126, miR-21 vs miR-10a, miR-10a vs miR-126, miR-126 vs miR-200c (Fig 4, A, B, D and F, respectively). Pathophysiology of T2DM is a complex of glycemic abnormality, damages of vascular endothelium, atherosclerosis, accumulation of extra vessel matrix. One miRNA has several hundreds of target genes and one gene has several binding sites for different miRNAs [16,17]. Thus, paired, or multiple miRNAs assay is expected to be more relevant to understand dysregulation of miRNAs than expression assay of a single miRNA [62]. Our data did not show any relationship between serum levels of miRNAs and laboratory data.

## Conclusion

Our study showed the limitation of the single miRNA analysis as a diagnostic tool of complicated pathophysiology of T2DM and suggested multi-miRNAs approach as the potential diagnostics for T2DM. In our study, 50 patients withT2DM and 15 control subjects were recruited. Further study with a larger cohort on usefulness of paired miRNAs analysis is recommended.

## Acknowledgment

We sincerely give many thanks to Hisae Nakashima, Tetsuhiro Harada, for their technical support to this study. This research was supported by JSPS KAKENHI Grant Number JP17K09020, 17H07059 and 20K08431.

